# mTORC1-mediated inhibition of 4EBP1 is essential for Hedgehog (HH) signaling and can be targeted to suppress HH-driven medulloblastoma

**DOI:** 10.1101/130872

**Authors:** Chang-Chih Wu, Shirui Hou, Brent A. Orr, Yong Ha Youn, Fanny Roth, Charles G. Eberhart, Young-Goo Han

**Affiliations:** Department of Developmental Neurobiology, 262 Danny Thomas Place, Memphis, TN 38105, USA; Department of Pathology, St. Jude Children’s Research Hospital, 262 Danny Thomas Place, Memphis, TN 38105, USA; Sorbonne Universités, UPMC Univ Paris 06, INSERM, CNRS, Centre de Recherche en Myologie (CRM), GH Pitié Salpêtrière, 47 bld de l’hôpital, Paris 13, France; Department of Pathology, The Johns Hopkins University School of Medicine, Ross Building 558, 720 Rutland Ave, Baltimore, MD 21205, USA

**Keywords:** Hedgehog, MTOR, 4EBP1, Medulloblastoma

## Abstract

Mechanistic target of rapamycin (MTOR) cooperates with Hedgehog (HH) signaling, but the underlying mechanisms are incompletely understood. Here, we provide genetic, biochemical, and pharmacologic evidence that MTOR complex 1 (mTORC1)-dependent translation is a prerequisite for HH signaling. The genetic loss of mTORC1 function inhibited HH signaling– driven growth of the cerebellum and medulloblastoma. Inhibiting translation or mTORC1 blocked HH signaling. Depleting 4EBP1, an mTORC1 target that inhibits translation, alleviated the dependence of HH signaling on mTORC1. Consistent with this, phosphorylated 4EBP1 levels were elevated in HH signaling–driven medulloblastomas in mice and humans. In mice, an mTORC1 inhibitor suppressed medulloblastoma driven by a mutant SMO that is resistant to an SMO inhibitor in the clinic, prolonging the survival of the mice. Our study reveals mTORC1-mediated translation to be a key component of HH signaling and an important target for treating medulloblastoma and other cancers driven by HH signaling.

## INTRODUCTION

Hedgehog (HH) signaling regulates many aspects of animal development from early embryonic patterning to tissue homeostasis in adults (Briscoe and Therond, 2013). Defective HH signaling causes various developmental malformations, whereas aberrantly active HH signaling can lead to tumors, including medulloblastomas, basal cell carcinomas, rhabdomyosarcomas, meningiomas, and odontogenic tumors (Amakye et al., 2013). Elucidating the molecular mechanism of HH signal transduction is critical for understanding normal development and diseases, including congenital defects and tumors arising from abnormal HH signaling.

In vertebrates, canonical HH signaling is triggered by one of three HH proteins (Sonic Hedgehog [SHH], Indian Hedgehog, or Desert Hedgehog) and culminates in changes in the transcriptional program via a series of inhibitory signaling cascades. In the absence of HH ligands, Patched1 (PTCH1), a 12-transmembrane receptor, localizes to the primary cilium and constitutively inhibits ciliary accumulation and the activation of a G protein–coupled receptor (GPCR)-like protein Smoothened (SMO) (Rohatgi et al., 2007). HH binds to and inhibits PTCH1, leading to ciliary accumulation and the activation of SMO (Corbit et al., 2005; Rohatgi et al., 2007). Activated SMO promotes the activation of GLI transcription factors (GLI2 and GLI3) by inhibiting their binding partner Suppressor of Fused (SUFU) and protein kinase A (PKA) (Hui and Angers, 2011). Activated GLI2 and GLI3 transcription factors induce the expression of HH target genes, including *Ptch1* and *Gli1*, which function, respectively, as a negative feedback regulator and a positive feed-forward regulator of HH signaling (Briscoe and Therond, 2013; Hui and Angers, 2011).

During perinatal development, granule neuron precursor cells (GNPs) in the cerebellum undergo massive proliferation to produce the cerebellar granule neurons that constitute more than half of the neurons in the brain. This proliferation of GNPs is driven by HH signaling (Dahmane and Ruiz i Altaba, 1999; Wallace, 1999; Wechsler-Reya and Scott, 1999), and germline and somatic mutations that lead to constitutive activation of HH signaling in GNPs result in unconstrained proliferation of GNPs and the development of SHH-subgroup medulloblastomas (Hahn et al., 1996; Johnson et al., 1996; Lam et al., 1999; Raffel et al., 1997; Reifenberger et al., 1998; Taylor et al., 2002; Vorechovsky et al., 1997; Wolter et al., 1997). Medulloblastoma is the most common pediatric brain cancer, and the SHH subgroup constitutes a third of these tumors. Aberrant activation of HH signaling in GNPs also causes medulloblastoma in mouse models (Oliver et al., 2005; Schuller et al., 2008; Yang et al., 2008). Accordingly, SMO inhibitors have demonstrated efficacy in treating SHH-subgroup medulloblastoma in both clinical trials and mouse models (Berman et al., 2002; Buonamici et al., 2010; Robinson et al., 2015; Rodon et al., 2014; Romer et al., 2004; Rudin et al., 2009). However, medulloblastoma cells inevitably acquire resistance to SMO inhibitors in humans and mice (Buonamici et al., 2010; Dijkgraaf et al., 2011; Robinson et al., 2015; Rudin et al., 2009; Yauch et al., 2009). Furthermore, tumors driven by alterations downstream from SMO do not respond to SMO inhibitors (Buonamici et al., 2010; Dijkgraaf et al., 2011; Robinson et al., 2015). Therefore, new therapies are needed to overcome these resistances and target tumors with mutations downstream from SMO.

Mechanistic target of rapamycin (MTOR) is a kinase that critically regulates the homeostasis and growth of cells and organisms in response to diverse extracellular and intracellular cues (Laplante and Sabatini, 2012). MTOR forms two multi-protein complexes, MTOR complex 1 (mTORC1) and mTORC2 (Figure S1A). mTORC1 is a key regulator of protein translation (Ma and Blenis, 2009). The two major downstream effectors of mTORC1 in translational control are ribosomal protein S6 (RPS6) kinase 1 (RPS6KB1, better known as S6K1) and eukaryotic translation initiation factor 4E (EIF4E) binding protein 1 (EIF4EBP1, better known as 4EBP1). 4EBP1 binds to EIF4E bound at the 5′ cap of mRNAs and prevents the formation of the translation initiation complex (Sonenberg and Hinnebusch, 2009). Phosphorylation of 4EBP1 by mTORC1 dissociates 4EBP1 from EIF4E, thus enabling the formation of the translation initiation complex. Deregulated protein translation is critical for the development of many cancers. Oncogenic signaling pathways often aberrantly increase EIF4E activity by enhancing its transcription and/or by inhibiting 4EBP1 through mTORC1-dependent phosphorylation (Truitt and Ruggero, 2016). HH signaling increases EIF4E transcription in GNPs (Mainwaring and Kenney, 2011); however, it is unclear whether HH signaling controls 5′ cap–dependent translation and whether oncogenic HH signaling depends on the mTORC1/4EBP1/EIF4E axis.

Here, we report that HH signaling promotes mTORC1/4EBP1-dependent translation and mTORC1/4EBP1-dependent translation is essential for HH signaling. Disrupting mTORC1 function inhibited GNP expansion during development and inhibited medulloblastoma growth driven by a mutant SMO that is resistant to an SMO inhibitor currently in clinical use. mTORC1 is essential for HH signaling and may be an important target for treating HH signaling–driven medulloblastoma and other cancers and for overcoming resistance to SMO inhibitors.

## RESULTS

### mTORC1 is required for HH signaling–driven growth of the cerebellum and medulloblastoma

Earlier studies concluded that HH signaling does not require mTOR because rapamycin (an allosteric inhibitor of MTOR) does not block HH signaling from inducing target-gene expression (Mainwaring and Kenney, 2011; Riobo et al., 2006). However, rapamycin only partially inhibits MTOR activities (Choo et al., 2008; Thoreen et al., 2009). Furthermore, MTOR cooperates with HH signaling in several types of cancer cell and contributes to the development of resistance to an SMO inhibitor (Buonamici et al., 2010; Filbin et al., 2013; Kern et al., 2015; Sharma et al., 2015; Syu et al., 2016; Wang et al., 2012). To scrutinize the function of MTOR in HH signaling *in vivo*, we conditionally removed Raptor and Rictor, essential components of mTORC1 and mTORC2, respectively (Figure S1A), in GNPs by using *GFAP::Cre* (Hara et al., 2002; Jacinto et al., 2004; Kim et al., 2002; Sarbassov et al., 2004). *GFAP::Cre* results in recombination in GNPs that require HH signaling for their proliferation (Spassky et al., 2008; Zhuo et al., 2001). The cerebella of *GFAP::Cre; Raptor*^*loxP/loxP*^ mice were much smaller than those of wild-type (WT) mice and contained far fewer granule neurons (Figure 1A). The small cerebellum containing fewer granule neurons is a stereotypical phenotype that results from defective HH signaling in GNPs due to the loss of *Gli2*, *Smo*, or ciliogenic genes (*Kif3a* and *Ift88*) (Chizhikov et al., 2007; Corrales et al., 2006; Spassky et al., 2008). Consistent with defective HH signaling, GNPs isolated from *GFAP::Cre; Raptor*^*loxP/loxP*^ mice had dramatically reduced levels of GLI1, a direct target of HH signaling, and SMO (Figure 1B). In contrast to the loss of Raptor, the loss of Rictor did not affect cerebellar growth in *GFAP::Cre; Rictor*^*loxP/loxP*^ mice (Figure 1A). Thus, mTORC1, but not mTORC2, was required for HH signaling to expand GNPs during cerebellar development *in vivo*.

**Figure 1.**
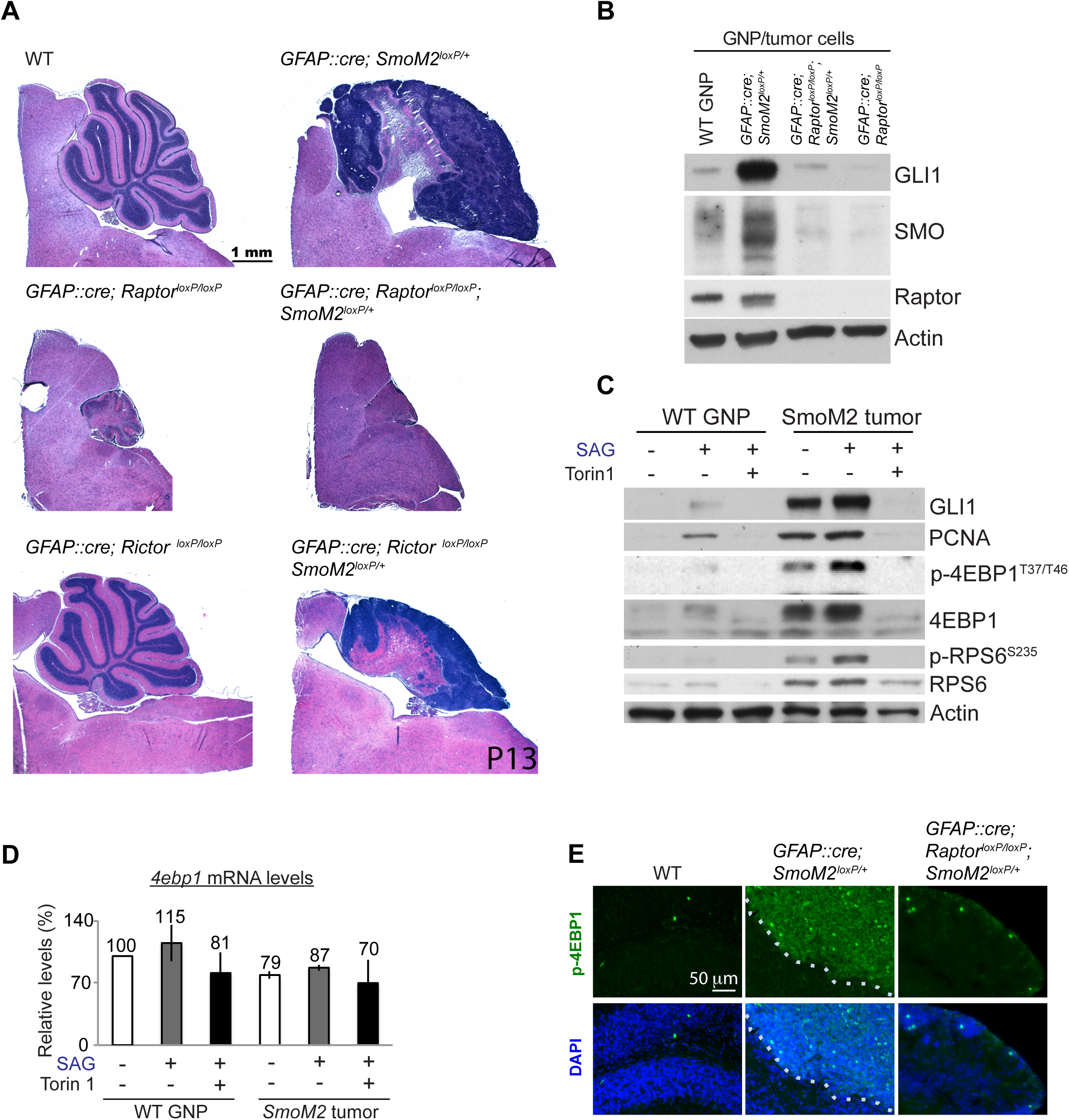
mTORC1 is required for HH-driven cerebellar and medulloblastoma development. (A) Hematoxylin and eosin–stained sagittal sections of the cerebellum. The loss of Raptor in GNPs inhibited cerebellar and medulloblastoma development. (B) The loss of Raptor decreased GLI1 and SMO protein levels in GNPs and in tumor cells purified from the cerebella of P7 WT and mutant mice. (C) Analysis of the indicated protein levels in WT GNPs and tumor cells isolated from P7 WT and *GFAP::Cre*; *SmoM2*^*loxP/+*^ mice, respectively. Isolated cells were grown in culture for 36 h with or without SAG and Torin1. Torin1 decreased the high levels of GLI1, SMO, 4EBP1, p-4EBP1, and PCNA in tumor cells and SAG-treated GNPs. (D) SAG and Torin1 treatment did not affect the *4ebp1* mRNA levels in GNPs or *SmoM2* tumor cells in culture. (E) Immunofluorescence labeling showing the high p-4EBP1 levels in *GFAP::Cre; SmoM2*^*loxP/+*^ mice, which were absent in WT or *GFAP::Cre; Raptor*^*loxP/loxP*^*; SmoM2*^*loxP/+*^ mice. Scattered mitotic cells were strongly labeled by the antibody to p-4EBP1.

Next, we asked whether mTORC1 is also required for oncogenic HH signaling. *GFAP::Cre; SmoM2*^*loxP/+*^ mice, which express in their GNPs an oncogenic mutant SMO (SMOM2) found in patients with basal carcinoma, medulloblastoma, or meningioma (Brastianos et al., 2013; Lam et al., 1999; Xie et al., 1998), develop medulloblastoma by postnatal day (P) 7 to 10 (Han et al., 2009). Remarkably, disrupting mTORC1 in *GFAP::Cre; SmoM2*^*loxP/+*^; *Raptor*^*loxP/loxP*^ mice prevented medulloblastoma development (Figure 1A). SMO and GLI1 levels were greatly reduced in GNPs isolated from *GFAP::Cre; SmoM2*^*loxP/+*^; *Raptor*^*loxP/loxP*^ mice (Figure 1B). In contrast, the loss of Rictor in *GFAP::Cre; SmoM2*^*loxP/+*^; *Rictor*^*loxP/loxP*^ mice did not prevent tumor development, although the tumors were smaller than those in *GFAP::Cre; SmoM2*^*loxP/+*^ mice (Figure 1A). Thus, mTORC1, but not mTORC2, was required for an oncogenic HH signaling pathway to induce medulloblastoma *in vivo*.

To directly examine the role of MTOR in HH signaling in GNPs and medulloblastoma cells, we treated primary WT GNPs and medulloblastoma cells with SAG (a SMO agonist) and Torin1 (an active-site mTOR inhibitor) in culture. Consistent with the *in vivo* phenotype, Torin1 dramatically decreased the SAG-or SMOM2-induced levels of GLI1 and PCNA (proliferating cell nuclear antigen, a proliferation marker) in WT GNPs and in *GFAP::Cre; SmoM2*^*loxP/+*^ medulloblastoma cells in primary culture (Figure 1C). Thus, MTOR was required for HH signaling in GNPs and in SMOM2-driven medulloblastoma cells to increase GLI1 and PCNA levels.

Notably, the total 4EBP1 and phosphorylated 4EBP1 (p-4EBP1) protein levels, but not the *4ebp1* mRNA levels, were dramatically increased in WT GNPs treated with SAG and in medulloblastoma cells (Figures 1C and 1D). Torin1 abolished these increases in 4EBP1 proteins without decreasing the *4EBP1* mRNA levels (Figures 1C and 1D). Together, these results suggest that HH signaling post-transcriptionally increases 4EBP1 levels, possibly by mTORC1-dependent translation, and increases their mTORC1-dependent phosphorylation. The level of p-4EBP1 was also greatly increased in medulloblastoma tissues of *GFAP::Cre; SmoM2*^*loxP/+*^ mice (Figure 1E), thereby demonstrating that there are strong mTORC1 activities in tumor cells *in situ*. The critical role of mTORC1 in medulloblastoma growth in G*FAP::Cre; SmoM2*^*loxP/+*^ mice may be to phosphorylate and antagonize the immensely increased levels of 4EBP1, a negative regulator of 5′ cap–dependent translation.

### MTOR/4EBP1-dependent translation is required for canonical HH signaling

To investigate the mechanism by which MTOR functions in HH signaling, we employed NIH 3T3 mouse embryonic fibroblast cells. NIH 3T3 cells have been used extensively to study the molecular and cellular mechanisms of HH signaling, because they contain components required for canonical HH signaling and produce robust and consistent responses to HH signaling. In NIH 3T3 cells, the activation of HH signaling by SAG increased the mRNA levels of *Gli1*, a direct transcriptional target of canonical HH signaling (Figure 2A). Torin1 (an active-site MTOR inhibitor), but not rapamycin (an allosteric MTOR inhibitor), completely suppressed this *Gli1* mRNA induction (Figure 2A). Thus, rapamycin-resistant MTOR activities were required downstream from SMO for canonical HH signaling. SMO must localize to cilia in order to activate canonical HH signaling (Corbit et al., 2005). Torin1 did not inhibit the ciliary accumulation of SMO after SAG stimulation (Figure S2), thereby ruling out a possible mechanism of mTORC1 function in canonical HH signaling.

**Figure 2.**
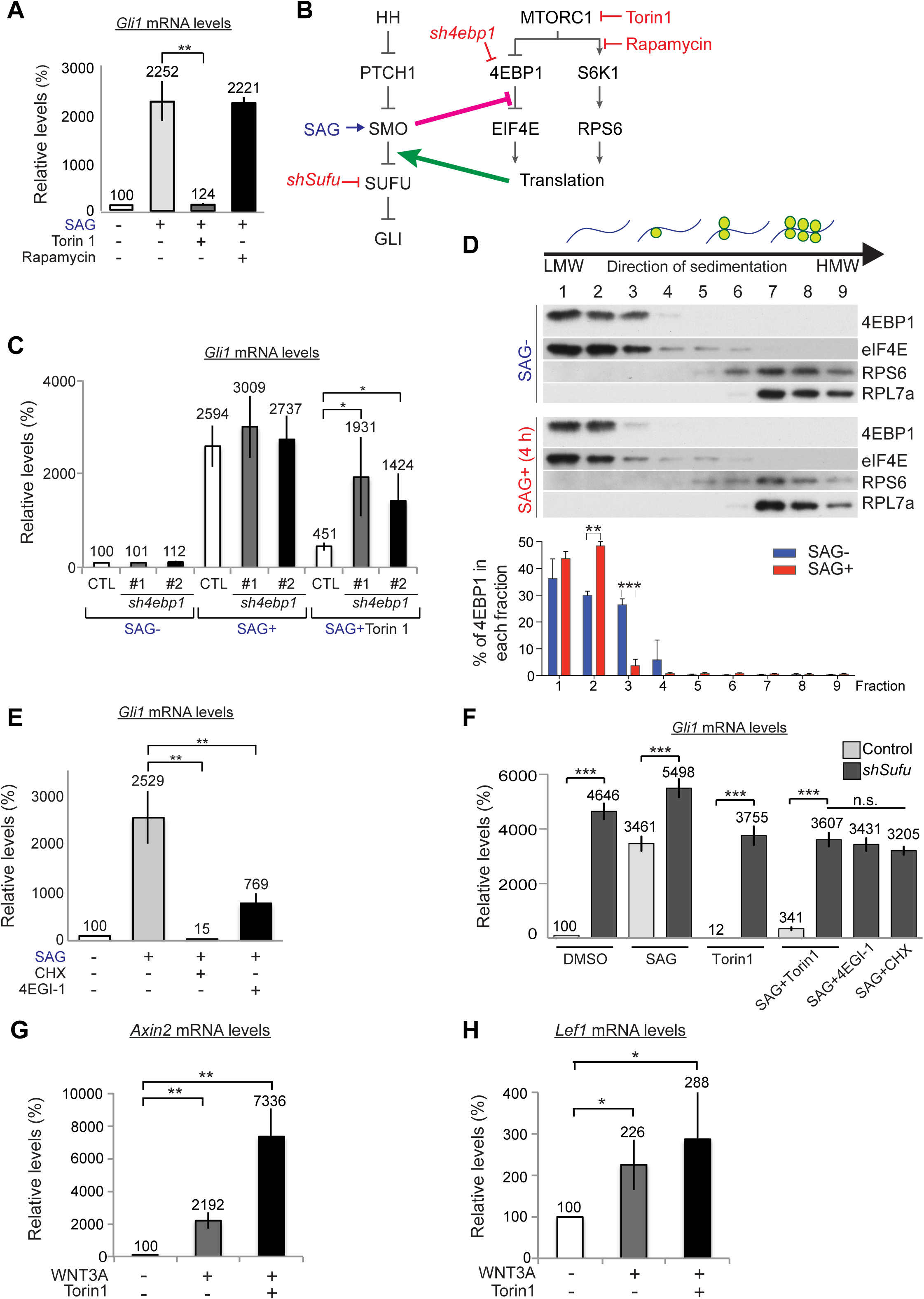
MTOR/4EBP1-dependent translation is required for HH signaling. (A) Torin1, but not rapamycin, suppressed the *Gli1* mRNA increase in SAG-stimulated NIH 3T3 cells. The bars show the fold changes in mRNA levels relative to those in untreated cells. (B) Simplified schematic of the HH and mTORC1 signaling cascade with the activators and inhibitors used in the study. We found that activation of SMO inhibits 4EBP1/EIF4E complex formation and that translation is required between SMO and SUFU in canonical HH signaling. (C) Depleting 4EBP1 restored the SAG-induced *Gli1* mRNA increase in Torin1-treated NIH 3T3 cells. (D) Western blotting and quantification (graphs) show that 4 h of SAG stimulation shifted 4EBP1 to the lighter cytosolic fractions in cell lysates fractionated on a 17%–47% sucrose gradient. Note the changes in 4EBP1 in fractions 2 and 3 of the SAG-treated NIH 3T3 cell lysates, as compared to the control cell lysates. The bars of the graph show the percentage of the total 4EBP1 in each fraction. The results are the average±SD of three independent experiments. The levels of ribosomal proteins (RPS6 and RPL7A) were used to determine the distribution of the cytosol, initiation complex, monosome, and polysomes, as shown in the schematic. (E) Translation inhibitors (CHX and 4EGI-1) suppressed the SAG-induced increase in *Gli1* mRNA levels in NIH 3T3 cells. (F) Depleting SUFU increased *Gli1* mRNA levels in the absence of SAG and in the presence of Torin1 or translation inhibitors. (G, H) Torin1 did not inhibit WNT3A from inducing the expression of the target genes *Axin2* (G) and *Lef1* (H). The bars in (A), (C), and (E) through (H) show the fold changes in mRNA levels relative to those in untreated control cells. The results shown in these panels are the average±SD of three independent experiments, each performed in triplicate. * *P<*0.05; ** *P<* 0.01; *** *P<* 0.005; n.s., *P*> 0.05.

S6K1 and 4EBP1 are two major substrates of mTORC1 in translational control (Figure 2B). Whereas Torin1 inhibits mTORC1 from phosphorylating both S6K1 and 4EBP1, rapamycin efficiently blocks S6K1 but not 4EBP1 phosphorylation (Choo et al., 2008; Thoreen et al., 2009), suggesting that inhibition of 4EBP1 by mTORC1-dependent phosphorylation is required for canonical HH signaling. Thus, we tested whether depletion of 4EBP1 can bypass the requirement of MTOR in canonical HH signaling. *4EBP1* knockdown (KD) by two different shRNAs did not increase *Gli1* mRNA levels in unstimulated cells, but it significantly restored the SAG-stimulated *Gli1* mRNA increase in the presence of Torin1 (Figure 2C), indicating that mTORC1-mediated inhibition of 4EBP1 is required for HH signaling.

Inhibitory phosphorylation of 4EBP1 by mTORC1 is a key step in the initiation of 5′ cap– dependent translation; phosphorylated 4EBP1 dissociates from EIF4E bound at the 5′ cap of the mRNA, thus enabling the formation of the translation initiation complex. To investigate whether HH signaling affected the complex of 4EBP1, EIF4E, and mRNA, we fractionated whole-cell lysates by using sucrose gradients and examined the distribution of key components of the translational machinery. SAG stimulation shifted 4EBP1 to the lighter cytosolic fractions within 4 h (compare the 4EBP1 levels in fractions 2 and 3 in Figure 2D), suggesting that HH signaling promotes translation by releasing 4EBP1 from the EIF4E-mRNA complex and that Torin1 inhibits canonical HH signaling by suppressing this HH-promoted translation. Indeed, like Torin1, inhibitors of translation initiation (4EGI-1) or elongation (cycloheximide [CHX]) suppressed *Gli1* mRNA induction by SAG (Figure 2E). Remarkably, however, mTORC1/4EBP1-dependent translation was dispensable for HH signaling triggered by the loss of SUFU, a key downstream negative regulator of HH signaling in mammals (Cooper et al., 2005; Svard et al., 2006) (Figure 2F, S1B, S1C). *Gli1* mRNA and GLI1 protein levels were strongly induced by *Sufu* knockdown in the absence of SAG or SHH (Figures 2F, S1B, S1C). *Sufu* KD cells still strongly increased *Gli1* mRNA and GL1 protein levels in the presence of Torin1 or translation inhibitors (4EGI-1 or CHX) (Figure 2F, S1B, S1C). Notably, 4EBP1 levels were increased by SAG but not by *Sufu* KD (Figure S1C) that induced *Gli1* mRNA expression, consistent with our finding that HH signaling increased 4EBP1 post-transcriptionally in GNPs (Figure 1C, 1D). Taken together, these results show that rapamycin-resistant and mTORC1/4EBP1-dependent translation was required for a step between SMO and SUFU in the canonical HH signaling pathway (Figure 2B).

### MTOR activities are not required for WNT signaling

MTOR might be a general prerequisite for signaling pathways. We asked whether MTOR was specifically required for HH signaling by testing its function in WNT signaling, which shares striking similarities with HH signaling (Kalderon, 2002). In both signaling pathways, common protein kinases (glycogen synthase kinase 3 and casein kinase 1) constitutively promote proteolytic processing of effector transcription factors (β-catenin for WNT and GLI2/3 for HH), and the activation of Frizzled-family GPCR-like proteins (Frizzled for WNT and SMO for HH) inhibits phosphorylation-dependent proteolytic processing of the effector transcription factors, leading to the accumulation and activation of β-catenin in WTN signaling and of GLI2/3 in HH signaling. In NIH 3T3 cells, WNT3A induced the expression of target genes, *Axin2* and *Lef1* (Filali et al., 2002; Hovanes et al., 2001; Jho et al., 2002; Lustig et al., 2002; Yan et al., 2001); however, blocking MTOR with Torin1 did not inhibit WNT signaling from inducing these WNT target genes but actually augmented their induction (Figures 2G and 2H). Thus, MTOR function is specifically required for HH signaling but not for WNT signaling, despite the similarities between the two.

Fibroblast growth factor 2 (FGF2) also increased *Gli1* mRNA levels in NIH 3T3 cells (Figure S1D). Although SANT1 (a SMO inhibitor) and Torin1 almost completely blocked SAG from increasing *Gli1* mRNA levels, they failed to efficiently block FGF2 from doing so (Figure S1D). Thus, in contrast to HH signaling, MTOR activities were not absolutely required for FGF2 to induce *Gli1* expression.

### HH signaling increases SMO protein expression via a noncanonical pathway

Because our data suggested that HH signaling promoted 5′ cap–dependent translation and translation was required for canonical HH signaling, we looked for a candidate protein that positively regulates HH signaling and whose translation is promoted by HH signaling. We noted that the loss of Raptor resulted in a dramatic decrease in SMO protein levels in the cerebellum (Figure 1B). *Smo* is not an established transcriptional target of HH signaling, suggesting that MTORC1 and HH signaling increase or maintain SMO protein levels. Consistent with these observations, activation of HH signaling via either SAG or SHH treatment (Figure 3A) increased SMO protein levels in both GNPs and NIH3T3 cells (Figures 3B–D). It is noteworthy that the increase in SMO occurred much earlier (at 8 h vs. 24 h) than the increase in GLI1, which is a direct transcriptional target of HH signaling (Figure 3C). In contrast, stimulating cells with fetal bovine serum (FBS), epidermal growth factor (EGF), or FGF2 did not increase SMO protein levels, although FBS and FGF2 increased GLI1 protein levels (Figures 3E and S3A), suggesting that the increase in SMO proteins was more specifically attributable to HH signaling activation than was the increase in GLI1. Consistently, SANT-1, a SMO antagonist, inhibited the SHH-induced increase in SMO and GLI1 proteins but not the FBS-induced increase in GLI1 proteins (Figure 3E).

**Figure 3.**
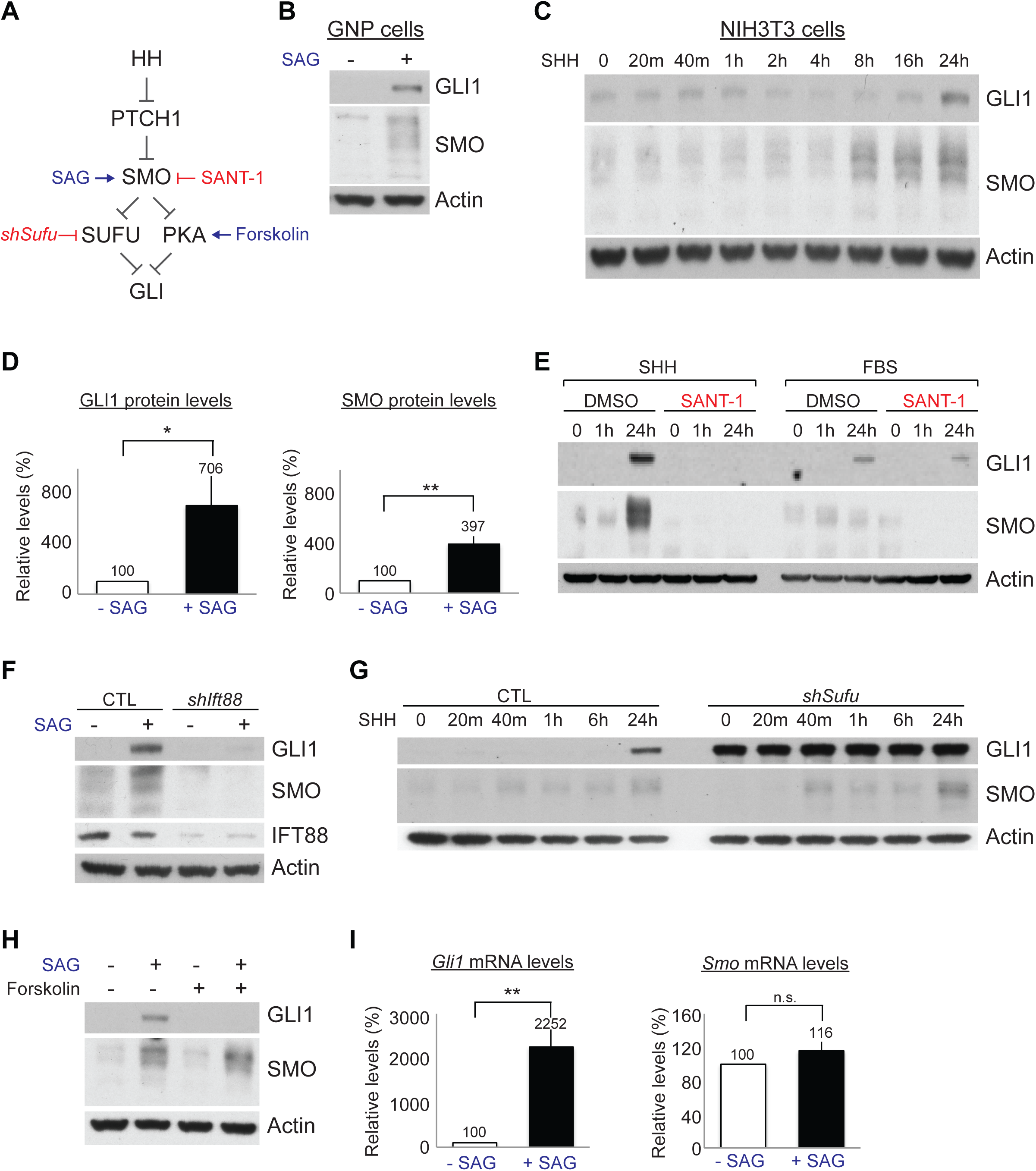
HH signaling increases SMO protein levels via a noncanonical pathway. (A) Schematic showing a simplified HH signaling pathway with reagents used to modulate the signaling components. (B) Western blot showing that 24 h of treatment with SAG, a SMO agonist, increased GLI1 and SMO protein levels in GNPs in culture. (C) GLI1 and SMO protein levels in NIH 3T3 cells at the indicated times after SHH treatment. Note that the SMO protein increased faster than the GLI1 protein. (D) Quantification of GLI1 and SMO protein levels 24 h after SAG treatment. The bars show the fold changes in the protein levels relative to those in untreated cells. The results are the average±SD of three independent experiments. (E) SMO and GLI1 protein levels in NIH 3T3 cells stimulated with SHH or FBS in the presence or absence of a SMO inhibitor, SANT-1. (F) SAG-induced GLI1 and SMO protein levels in NIH 3T3 cells were suppressed in cells expressing shRNA against *Ift88* (*shIft88*). (G) GLI1 and SMO protein levels in NIH 3T3 cells expressing *shSufu* at the indicated times after SHH treatment. Note that SMO, but not GLI1, was induced by SHH. (H) Forskolin blocked SAG from increasing the level of GL1 but not from increasing the level of SMO protein. (I) SAG increased the level of *Gli1* mRNA but not that of *Smo* mRNA in NIH 3T3 cells. The bars show the fold changes in the mRNA levels relative to those in untreated cells. The results are the average±SD of three independent experiments, each performed in triplicate. Actin was used as the loading control for Western blot experiments. * *P<*0.05; ** *P<* 0.01; n.s., *P*> 0.05.

HH-induced SMO protein increase preceded the increase of GLI1 (Figure 3C). Therefore, we examined whether central components of canonical HH signaling are required for the increase in SMO protein levels (Figure 3A). Primary cilia are essential for canonical HH signaling in mammals (Goetz and Anderson, 2010). The KD of *Intraflagellar transport 88* (*Ift88*), an essential ciliogenic gene (Pazour et al., 2000), blocked SAG from inducing the GLI1 and SMO proteins (Figure 3F). Thus, HH signaling required primary cilia in order to increase SMO protein levels. Similar to SANT-1 treatment (Figure 3E), *Ift88* KD lowered the basal SMO protein levels (lane 1 vs. 3 in Figure 3F), suggesting that basal HH signaling activity maintains SMO protein levels. Depleting SUFU, a key downstream negative regulator, strongly increased GLI1 levels in the absence of SHH or SAG (Figures 3G, S1B, S1C). However, SMO protein levels were increased only after SHH treatment in the *Sufu* KD cells (Figure 3G). PKA is another negative regulator of canonical HH signaling (Wang et al., 2000; Wang et al., 1999). The activation of PKA by forskolin completely blocked the increase in GLI1 protein levels after SAG stimulation but not the increase in SMO protein levels (Figure 3H). Canonical HH signaling culminates in changes in gene expression; however, *Smo* mRNA levels were not increased by SAG, whereas *Gli1* mRNA levels were increased (Figure 3I). Furthermore, *Ift88* KD diminished *Gli1* mRNA induction by SAG but did not affect *Smo* mRNA levels (Figures S3B, S3C), although it blocked SAG from increasing the SMO protein levels (Figure 3F). Taken together, these results demonstrate that HH signaling increased the SMO protein levels post-transcriptionally via a SUFU- and PKA-independent but SMO- and primary cilia–dependent pathway.

### HH signaling increases SMO translation in an mTORC1- and 4EBP1-dependent manner

Previous studies had shown that Hh signaling stabilizes Smo proteins in *Drosophila* (Alcedo et al., 2000; Denef et al., 2000). To test whether HH signaling in mammals also increased SMO by inhibiting its degradation, we asked whether interrupting the proteasome and lysosome functions, the two major mechanisms that degrade cellular proteins, increased SMO protein levels. SMO did not accumulate 8 h after we blocked proteasomes (with MG132 or bortezomib), lysosomes (with chloroquine), or both (with MG132 and chloroquine), by which time HH signaling had already increased the levels of SMO proteins (Figure S3D). In contrast, inhibiting translation initiation (with 4EGI-1) or elongation (with cycloheximide [CHX]) completely abolished SAG-induced SMO upregulation (Figure 4A), indicating that SMO upregulation is dependent on translation but not on increased stability. Moreover, Torin1, but not rapamycin, blocked the SAG-induced increase in SMO (Figure 4A). Torin1 inhibited SAG induction of SMO proteins in a dose-dependent manner but did not affect *Smo* mRNA levels (Figures 4B, 4C). To test whether Torin1 decreased SMO protein levels by destabilizing the protein, we inhibited the proteasomes simultaneously with the Torin1 treatment. Proteasome inhibition did not rescue the decrease in SMO proteins caused by Torin1 (Figure S3E). Taken together, these results show that HH signaling increased SMO protein synthesis and required rapamycin-resistant MTOR function in order to do so.

**Figure 4.**
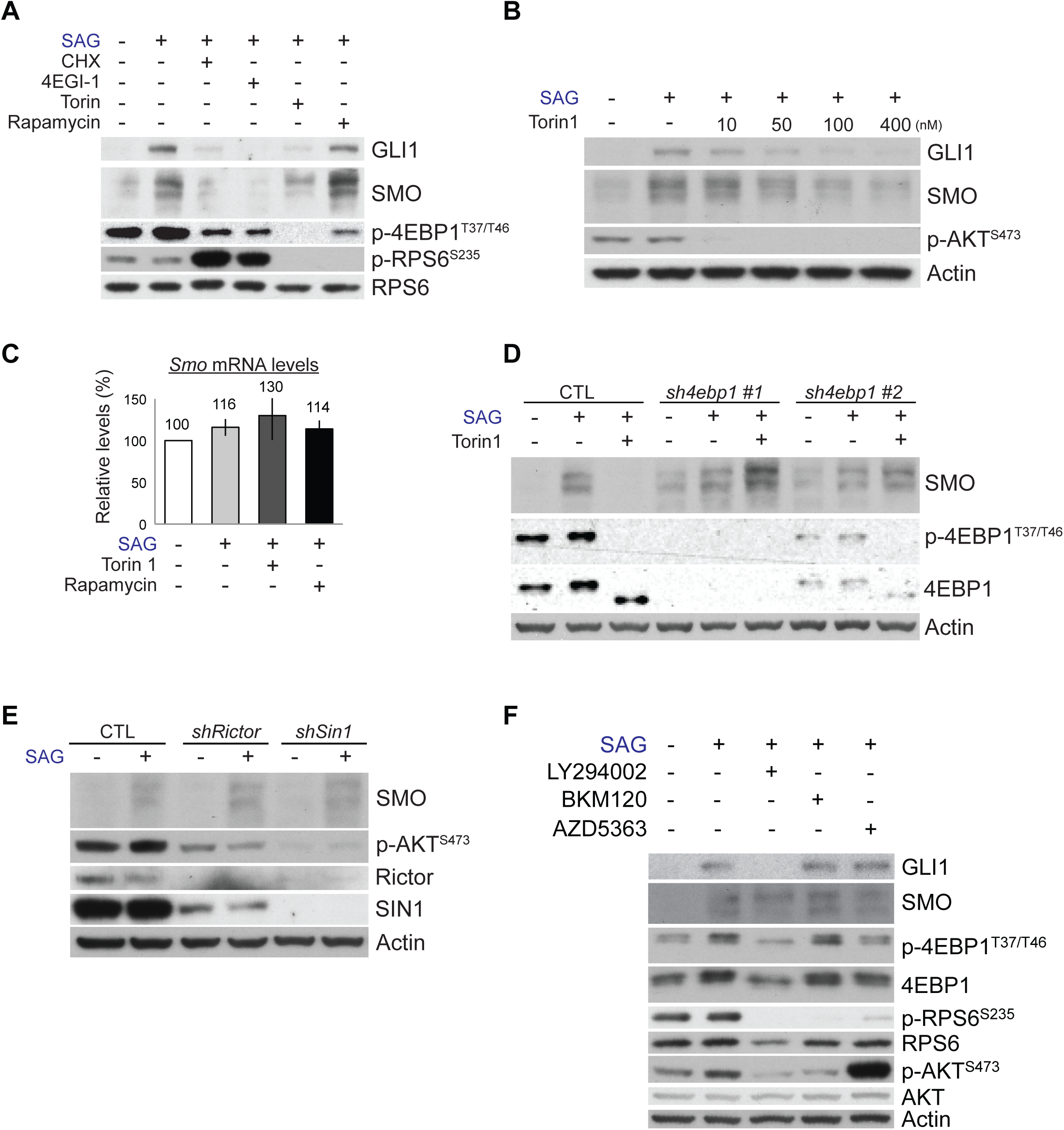
HH signaling increases SMO translation through an mTORC1- and 4EBP1-dependent mechanism. (A) Translation inhibitors (CHX, 4EGI-1) and Torin1, but not rapamycin, blocked the SAG-driven increase in SMO protein in NIH 3T3 cells. Note that Torin1, but not rapamycin, completely inhibited the phosphorylation of 4EBP1. (B) Torin1 suppressed SAG-induced GLI1 and SMO protein levels in a dose-dependent manner. (C) SAG, Torin1, and rapamycin did not affect *Smo* mRNA levels. The bars show the fold changes in the mRNA levels relative to those in untreated cells. The results are the average±SD of three independent experiments, each performed in triplicate. (D) Depleting 4EBP1 with two different shRNAs (*sh4ebp1* #1 and #2) increased the basal SMO protein levels and abolished the inhibitory effect of Torin1 on SMO induction. (E) Disruption of mTORC2 by shRNAs against Rictor (*shRictor*) or SIN1 (*shSin1*) decreased AKT phosphorylation but not SMO levels in SAG-treated NIH 3T3 cells. (F) PI3K inhibitors (LY294002, BKM120) or an AKT inhibitor (AZD5363) decreased p-RPS6 but not SAG-induced SMO protein levels in NIH 3T3 cells. The high level of p-AKT S473 in cells treated with AZD5363 represents feedback hyperphosphorylation after the inhibition of AKT.

The differential effects of rapamycin and Torin1 on SMO protein levels mirrored their differential effects on canonical HH signaling (Figure 2A) and suggested that inhibitory phosphorylation of 4EBP1 is required for HH signaling to increase SMO translation. Consistent with this, depleting 4EBP1 increased the basal SMO protein levels and restored SMO induction by SAG in the presence of Torin1 (Figure 4D). Thus, mTORC1-dependent inhibition of 4EBP1 was required for HH signaling to increase SMO protein synthesis.

Torin1, but not rapamycin, also inhibits mTORC2 (Thoreen et al., 2009). To test whether mTORC2 function is also required in order for HH signaling to increase SMO protein levels, we depleted either Rictor or MAPKAP1 (also known as SIN1), two essential components of mTORC2 (Jacinto et al., 2006; Jacinto et al., 2004; Sarbassov et al., 2004; Yang et al., 2006), in NIH 3T3 cells by using shRNAs. Depleting *Rictor* or *Sin1* decreased the phosphorylation of AKT serine/threonine kinase (AKT) at Ser473, a direct substrate of mTORC2 (Sarbassov et al., 2005), but failed to inhibit SAG from increasing SMO (Figure 4E). mTORC2 function was dispensable for the ability of HH signaling to increase SMO levels, similar to how it is dispensable in HH-driven cerebellar and medulloblastoma growth (Figure 1A).

Phosphatidylinositol 3-kinase (PI3K) and AKT are well-known upstream activators of mTORC1 (Laplante and Sabatini, 2012; Ma and Blenis, 2009) (Figure S1A); however, inhibitors of PI3K (LY294002 and BKM120) or AKT (AZD5363) failed to inhibit 4EBP1 phosphorylation or the SAG-induced increase in SMO protein (Figure 4F), although they efficiently inhibited the phosphorylation of RPS6. The high level of p-AKT S473 in cells treated with AZD5363 represents feedback hyperphosphorylation after the inhibition of AKT. LY294002 decreased GLI1 levels but not SMO levels. This decrease in GLI1 levels may result from PI3K-independent action of LY294002, because LY294002 can affect proteins unrelated to PI3K (Gharbi et al., 2007) and BKM120, a more selective PI3K inhibitor, did not affect GLI1 and SMO levels. A previous study also showed that another highly specific PI3K inhibitor (GNE-317) does not suppress HH signaling (Metcalfe et al., 2013). Taken together, these results show that mTORC1, but not mTORC2, PI3K, or AKT, was required for HH signaling to increase SMO translation.

### p-4EBP1 levels are high in SHH-subgroup medulloblastoma in humans

Our data showed that inhibitory phosphorylation of 4EBP1 by MTORC1 is essential for HH signaling. Consistent with this, SMOM2-driven mouse medulloblastoma was dependent on MTORC1 and had high levels of p-4EBP1 (Figure 1A, 1E). Thus, we examined whether p-4EBP1 levels were also high in human medulloblastoma. In humans, medulloblastomas are categorized in four subgroups: SHH, WNT, subgroup 3, and subgroup 4. The subgroups have distinct characteristics, including gene expression profiles, mutations, epigenomes, and signaling pathways that promote cancer (Taylor et al., 2012). Consistent with our mouse model, most SHH-subgroup human medulloblastomas (76%) expressed p-4EBP1 (Figure 5). Notably, p-4EBP1 staining was significantly enriched in the SHH subgroup as compared with non-WNT/non-SHH medulloblastomas (*P*=0.041 by the chi-square test), and the staining level in the SHH-subgroup medulloblastomas was greater than in the non-WNT/non-SHH subgroup tumors (*P*=0.020 by the Kruskal-Wallis test) (Figure 5B, S4), suggesting that there are strong mTORC1 activities in SHH-subgroup medulloblastomas.

**Figure 5.**
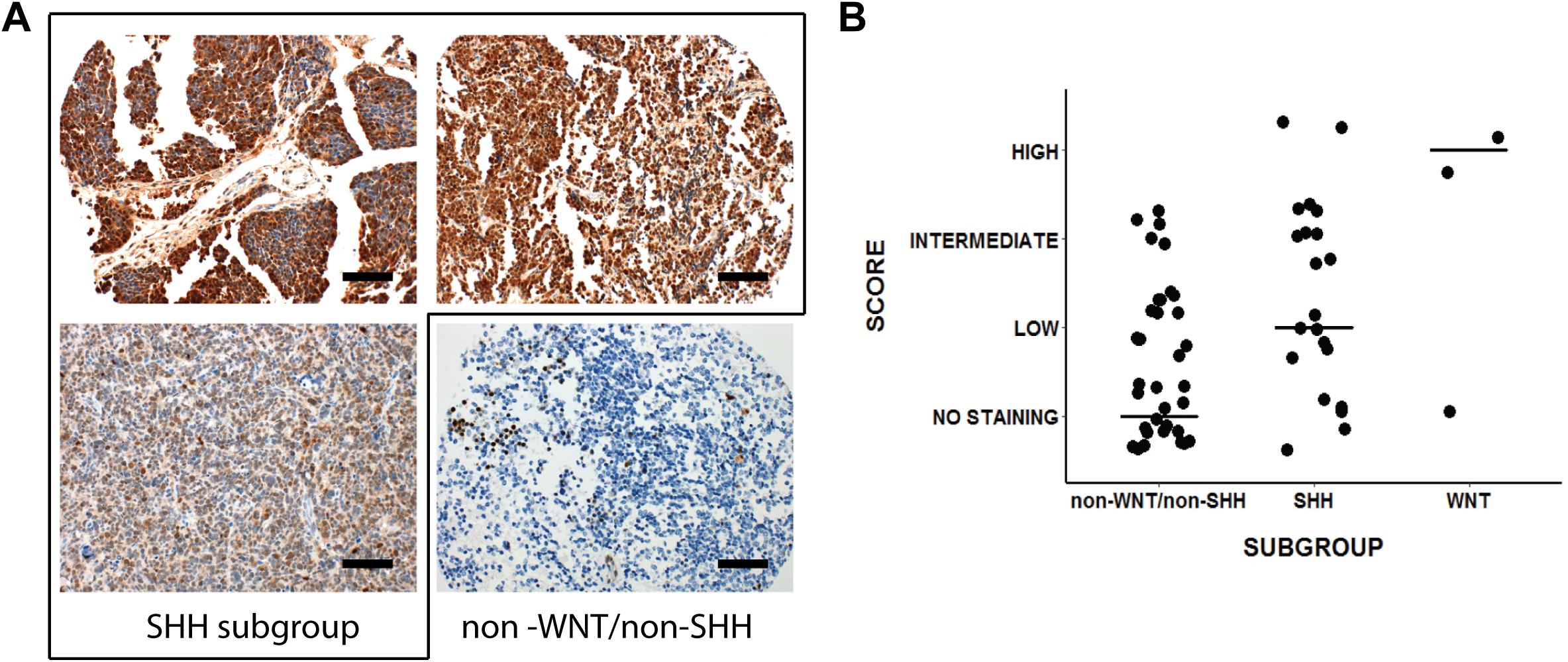
p-4EBP1 levels in human medulloblastoma. (A) Immunohistochemical staining for p-4EBP1 in human SHH-activated medulloblastoma samples demonstrated high-level immunoreactivity (top panels) or intermediate-level staining (bottom-left panel), whereas only focal, low-level staining was observed in non-WNT/non-SHH medulloblastoma (bottom-right panel). (B) The SHH-activated group exhibited higher levels of immunoreactivity compared to non-WNT/non-SHH tumors (*P<*=0.020 by the Kruskal-Wallis test). The graph has been jittered to enable visualization. The median staining level is represented by a line for each subgroup. Scale bar=200 μm.

### INK128, an MTOR inhibitor, penetrates the blood-brain barrier and inhibits HH signaling

The unexpected critical role of MTOR-dependent translation in HH signaling and the high p-4EBP1 levels in SHH-subgroup medulloblastoma in humans prompted us to investigate the potential of MTOR inhibitors in treating SHH-subgroup medulloblastoma. We chose INK128 (Hsieh et al., 2012) from among the commercially available MTOR inhibitors and tested whether it could penetrate the blood-brain barrier and inhibit HH signaling *in vivo*; INK128 is currently undergoing a clinical trial (NCT02133183) to study its brain penetrance and its effects on glioblastoma. Like Torin1, INK128 suppressed the SAG-induced increase in GLI1 and SMO in primary GNPs (Figure 6A). *In vivo*, INK128 inhibited the phosphorylation of mTORC1 substrates (RSP6 and 4EBP1) and an mTORC2 substrate (AKT) in WT cerebella within 2 h of administration by oral gavage at P10 (Figure 6B). The GLI1 and SMO levels were also decreased within 6 h after treatment (Figure 6B).

**Figure 6.**
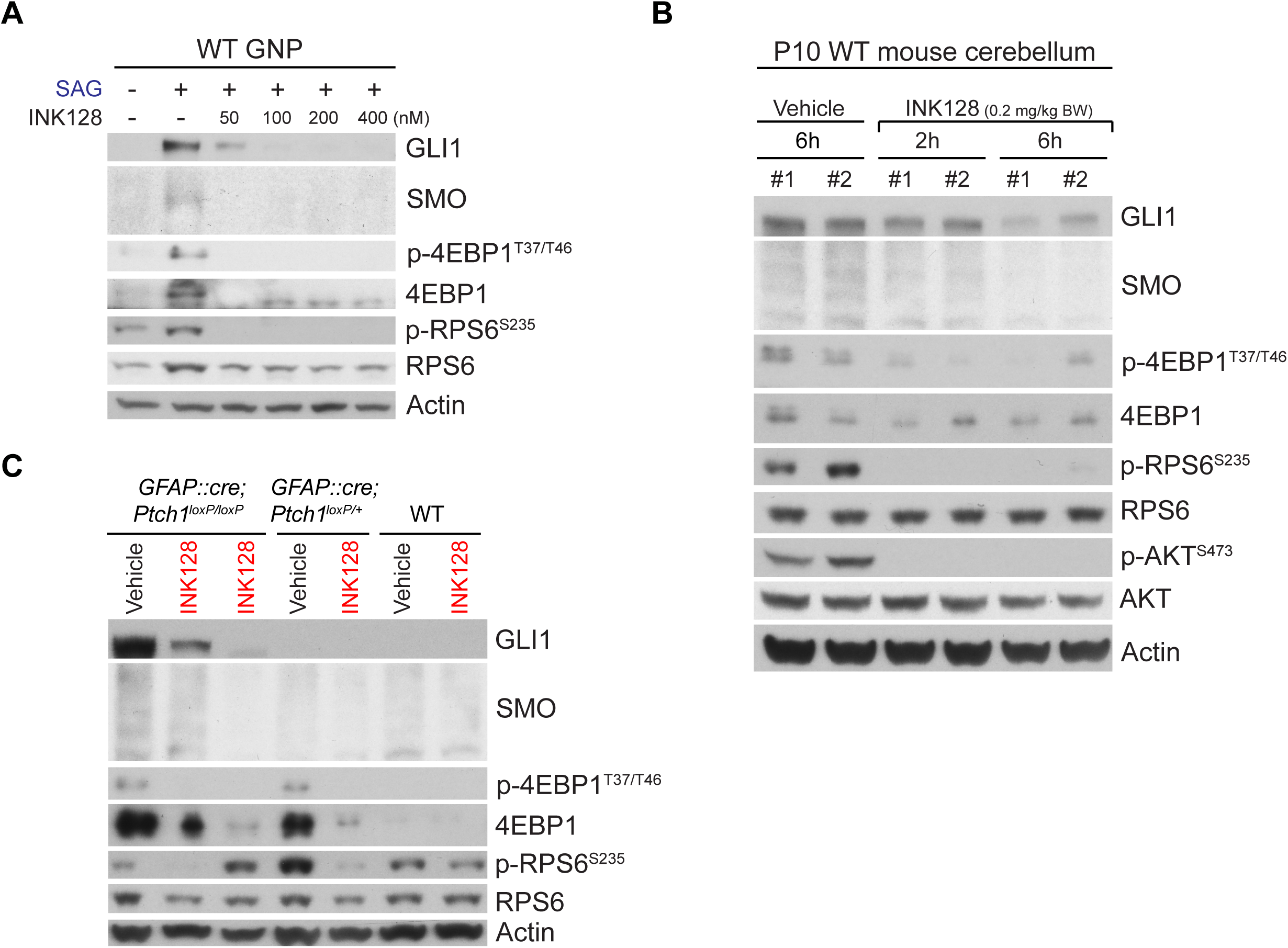
INK128 penetrates the brain and inhibits HH signaling. (A) INK128 suppressed the SAG-induced increase in GLI1, SMO, 4EBP1, and p-4EBP1 levels in GNPs in culture. (B) Administration of INK128 (0.2 mg/kg body weight [BW]) to WT pups by oral gavage at P10 inhibited the phosphorylation of 4EBP1, RPS6, and AKT within 2 h and decreased the GLI1 and SMO levels within 6 h. (C) Administration of INK128 (0.2 mg/kg BW) at P10 to WT, *GFAP::Cre; Ptch1*^*loxP/+*^, and *GFAP::Cre; Ptch1*^*loxP/loxP*^ mice decreased the high levels of GLI1, SMO, 4EBP1, and p-4EBP1 in the tumors of *GFAP::Cre; Ptch1*^*loxP/loxP*^ mice and the high levels of 4EBP1 and p-4EBP1 in the cerebella of *GFAP::Cre; Ptch1*^*loxP/+*^ mice by 24h. Mice treated only with vehicle were used as controls. Each lane in panels B and C represents an individual mouse. Actin was used as the loading control.

Next, we tested whether INK128 could inhibit HH signaling in medulloblastoma using *GFAP::Cre; Ptch1*^*loxP/loxP*^ mice, in which the loss of a tumor suppressor, PTCH1, results in greatly expanded GNPs by P2 and cerebella filled with tumor cells by P14 (Yang et al., 2008). Remarkably, a single dose of INK128 strongly suppressed the high levels of GLI1 and SMO in medulloblastoma in *GFAP::Cre; Ptch1*^*loxP/loxP*^ mice at 24 h after treatment at P10 (Figure 6C). 4EBP1 and p-4EBP1 levels were also higher in *GFAP::Cre; Ptch1*^*loxP/loxP*^ tumors than in WT cerebella, and they were suppressed by INK128. Thus, INK128 rapidly penetrated the blood-brain barrier and efficiently inhibited MTOR activities and HH signaling in WT cerebella and medulloblastomas. Notably, *GFAP::Cre; Ptch1*^*loxP/+*^ mice, which did not develop tumors and had levels of GLI1 and SMO comparable to those of WT mice, had much higher levels of 4EBP1 and p-4EBP1 than did WT mice, suggesting that 4EBP1 and p-4EBP1 levels are highly sensitive to HH signaling activities in GNPs.

### INK128 inhibits primary medulloblastoma growth in *GFAP::Cre; SmoM2^loxP/+^* mice and prolongs their survival

Because INK128 inhibited MTOR and HH signaling in medulloblastoma in *GFAP::Cre; Ptch1*^*loxP/loxP*^ mice, we next investigated whether the compound could suppress medulloblastoma growth driven by a mutant SMO (SMOM2) that is resistant to an SMO inhibitor in the clinic (Dijkgraaf et al., 2011). Currently, no targeted therapy is available for medulloblastoma that is resistant to SMO inhibitors, and an effective therapy for these tumors is urgently needed. *In vitro*, MTOR inhibition suppressed HH signaling in SMOM2-driven medulloblastoma cells isolated from *GFAP::Cre; SmoM2*^*loxP/+*^ mice (Figure 1C).

INK128 suppressed MTOR activities in *GFAP::Cre; SmoM2*^*loxP/+*^ medulloblastoma in a dose-dependent manner within 2 h of administration at P10 (Figure 7A). Note that the 4EBP1 and p-4EBP1 levels were dramatically increased in *GFAP::Cre; SmoM2*^*loxP/+*^ medulloblastoma but were suppressed by INK128 (Figure 7A), consistent with observations made *in vitro* (Figures 1C and 6A), in immunostained tumors (Figure 1E, 5A), and in *GFAP::Cre; Ptch1*^*loxP/loxP*^ mice (Figure 6C). Two days of daily treatment with INK128 decreased the number of tumor cells and the levels of GLI1 and the proliferation markers Ki67 and PCNA, as compared to the levels in mice treated with vehicle alone (Figures 7B, 7C). Daily treatment of INK128 for 8 days decreased tumor size; however, administering the same treatments to WT mice did not affect the size or histologic appearance of their brains (Figure 7D). All but one *GFAP::Cre; SmoM2*^*loxP/+*^ mouse treated with vehicle died within 18 days of birth (Figure 7E). Remarkably, daily administration of INK128 to *GFAP::Cre; SmoM2*^*loxP/+*^ mice, starting from P7 (when their entire cerebellum is essentially tumor tissue), significantly prolonged their survival (Figure 7E). Thus, inhibition of MTOR suppressed HH-driven medulloblastoma growth and may be effective against tumors driven by mutant SMO proteins that are resistant to SMO inhibitors.

**Figure 7.**
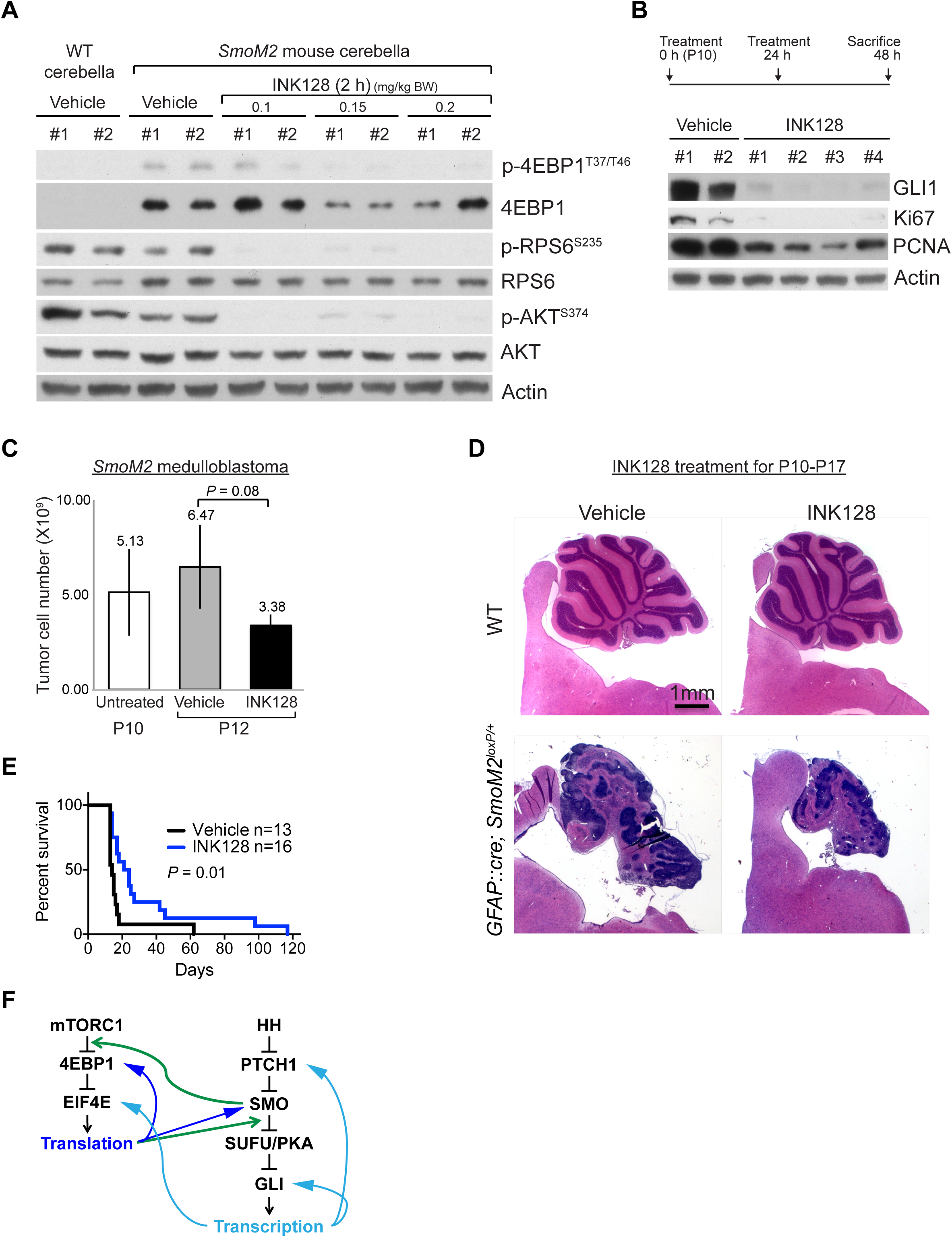
INK128 inhibits primary medulloblastoma growth in *GFAP::Cre; SmoM2^loxP/+^* mice and prolongs their survival. (A) Oral gavage of INK128 at P10 decreased the phosphorylation of 4EBP1, RPS6, and AKT in medulloblastomas of *GFAP::Cre; SmoM2*^*loxP/+*^ mice. Note the high levels of 4EBP1 and p-4EBP1 in vehicle-treated tumors compared with those in the WT cerebella at P10. (B, C) There was a large decrease in the levels of GLI1 and proliferation markers (Ki67 and PCNA) (B) and in the number of tumor cells (C) after 2 days of daily treatment with INK128 (0.2 mg/kg BW). (D) Hematoxylin and eosin–stained sagittal sections of cerebella showing the decrease in tumor size after 8 days of daily treatment with INK128 (0.2 mg/kg BW). The same treatments did not affect the size or histologic appearance of WT cerebella. (E) Daily treatments of INK128 (0.2 mg/kg BW) from P7 significantly prolonged the survival of *GFAP::Cre; SmoM2*^*loxP/+*^ mice, all but one of which died by P18 when treated with vehicle. (F) Schematic summary of crosstalk between HH and mTORC1 pathways. Activation of SMO releases the inhibitory association of EIF4E and 4EBP1 in an mTORC1-dependent manner, leading to increased translation of proteins, including SMO and 4EBP1. mTORC1/4EBP1-dependent translation is required as a step between SMO and SUFU in HH signaling to induce the transcription of target genes, including *Ptch1*, *Gli1*, and *Eif4e*.

## DISCUSSION

Canonical HH signaling activates transcription through a PTCH1-SMO-SUFU/PKA-GLI– dependent pathway. Here, we have shown that mTORC1/4EBP1-dependent translation is required for a step between SMO and SUFU in canonical HH signaling (Figure 7F). HH signaling induced dissociation of 4EBP1 from EIF4E-mRNA complex, promoting translation. HH signaling rapidly promoted *SMO* mRNA translation through a noncanonical, SUFU/PKA-GLI–independent, and mTORC1/4EBP1–dependent pathway. Translation inhibitors and MTOR inhibitors prevented HH signaling from inducing not only SMO protein synthesis but also *Gli1* transcription. Remarkably, MTOR activities were not required in order for HH signaling to induce transcription or translation in cells in which 4EBP1 was depleted. Thus, inhibitory phosphorylation of 4EBP1 by mTORC1 is essential for HH signaling to activate both translation and transcription. In contrast, MTOR activities were not required for WNT signaling, which is closely related to HH signaling.

Our study reveals mTORC1 to be a potentially important molecular target for treating SHH-subgroup medulloblastomas, most of which exhibited elevated levels of p-4EBP1. The genetic disruption of mTORC1 function prevented the development of medulloblastoma in *GFAP::Cre; SmoM2*^*loxP/+*^; *Raptor*^*loxP/loxP*^ mice. Moreover, INK128, an active-site MTOR inhibitor, readily penetrated the blood-brain barrier, strongly suppressed medulloblastoma growth, and significantly prolonged the survival of *GFAP::Cre; SmoM2*^*loxP/+*^ mice. It is noteworthy that medulloblastomas developed fully in young *GFAP::Cre; SmoM2*^*loxP/+*^ pups at a developmental stage similar to that at which medulloblastoma develops in human patients and that INK128 treatments were tolerated by and effective in pups whose cerebella consisted mostly of tumor tissue. Current therapies for medulloblastoma fail to cure approximately 20% to 30% of patients, and children who do survive the tumor suffer lifelong debilitating side effects from the treatments. Therefore, new therapies are needed that directly target the mechanisms that drive medulloblastoma. Until now, SMO has been the only molecule targeted in clinical trials of medulloblastoma treatments. Despite their promising efficacy, SMO inhibitors have shown some limitations in clinical trials; patients with mutations downstream from SMO do not respond to SMO inhibitors, and initial responders inevitably acquire resistance (Robinson et al., 2015; Rodon et al., 2014; Rudin et al., 2009). Our results show that MTOR inhibitors can overcome some of these limitations; thus, targeting MTOR in combination with the use of SMO inhibitors and/or other anti-cancer agents may be a promising strategy to treat SHH-subgroup medulloblastoma and other cancers driven by aberrant HH signaling.

PI3K and AKT are well-known upstream activators of mTORC1 (Laplante and Sabatini, 2012; Ma and Blenis, 2009). Mutations activating PI3K signaling have been found in SHH-subgroup medulloblastoma (Northcott et al., 2012; Robinson et al., 2012). Activation of PI3K signaling is a potential mechanism for the emergence of resistance to SMO inhibitors, and co-inhibition of SMO and PI3K blocks the phosphorylation of RPS6 and significantly delayed the appearance of resistance to SMO inhibitors in a mouse model (Buonamici et al., 2010; Metcalfe et al., 2013). Importantly, a derivative of rapamycin also delayed the development of resistance in a similar manner to PI3K inhibitors (Buonamici et al., 2010), suggesting that a rapamycin-sensitive mTORC1 function is important for resistance development. However, our results suggest that the function of mTORC1 in HH signaling is independent of its role in the development of resistance to SMO inhibitors; PI3K and AKT inhibitors and rapamycin, all of which blocked phosphorylation of RPS6 but not of 4EBP1, failed to inhibit HH signaling. A previous study also showed that a PI3K inhibitor (GNE-317) does not suppress HH signaling (Metcalfe et al., 2013). These results are consistent with the inefficiency of PI3K inhibitors, when administered as a single agent, at suppressing medulloblastoma growth (Buonamici et al., 2010; Metcalfe et al., 2013). In contrast, INK128 strongly suppressed HH signaling and the growth of medulloblastoma. Thus, active-site MTOR inhibitors may offer the advantage of directly inhibiting HH signaling, in addition to inhibiting the rapamycin-sensitive MTOR activities that contribute to resistance to SMO inhibitors.

HH signaling promotes transcription by relieving repression and regulates its activities by inducing the expression of positive- and negative-feedback regulators, namely GLI1 and PTCH1. Translational control by HH signaling shows striking similarities to transcriptional control; it promotes translation by relieving the inhibitory function of 4EBP1 and induces the expression of key positive and negative regulators of 5′ cap–dependent translation, namely EIF4E and 4EBP1, respectively (this study and Mainwaring and Kenney, 2011). HH signaling also promotes internal ribosome entry site–dependent translation of ornithine decarboxylase (ODC), and an ODC inhibitor suppresses the growth of SHH-subgroup medulloblastoma allografts (D’Amico et al., 2015). A recent study showed that HH signaling guides axons via inducing local translation at the growth cone (Lepelletier et al., 2017). Together, these results indicate that HH signaling–regulated translation constitutes critical parts of the HH signaling network in development and tumorigenesis. Deregulated translation control is a hallmark of human cancers and is critical for tumorigenesis downstream from multiple oncogenic signaling pathways (Truitt and Ruggero, 2016). Identifying mRNAs whose translation is regulated by HH signaling will deepen our understanding of HH signaling mechanisms and reveal potential therapeutic targets for HH-driven cancers.

## EXPERIMENTAL PROCEDURES

### Cell culture and stimulation

NIH 3T3 cells were grown in culture in DMEM containing 10% calf serum (Sigma) and 1× penicillin-streptomycin-glutamine (PSG; ThermoFisher). Cells were seeded at 1 million cells per 10-cm dish and typically reached 100% confluency 48 h after seeding. For serum starvation, confluent cells were maintained in DMEM containing 0.5% calf serum and 1× PSG. After 24 h of serum starvation, the cells were treated with the following agents as indicated in the text and figures: SHH peptides (200 ng/mL; R&D Systems), SAG (200nM; Cayman Chemical, catalog no. 11914), fetal bovine serum (10%; ThermoFisher), FGF2 (100 ng/mL; PeproTech), EGF (100 ng/mL; PeproTech), WNT3A (100 ng/mL; R&D Systems), SANT-1 (100 nM; Santa Cruz Biotechnology), Torin1 (400 nM; Tocris Bioscience), INK128 (400 nM; Selleck Chemicals), rapamycin (100 nM; LC Laboratories), BKM120 (400 nM; Selleck Chemicals), LY294002 (10 μM; LC Laboratories), AZD5363 (100 nM; Selleck Chemicals), 4EGI-1 (10 μM; EMD Millipore), cycloheximide (CHX) (100 μM; Sigma-Aldrich), forskolin (10 μM; Selleck Chemicals), MG132 (10 μM; Sigma-Aldrich), chloroquine (10 μM; Sigma-Aldrich), or Bortezomib (100 nM; Selleck Chemicals). All treatments were done for 24 h except those indicated otherwise sin the text and figures. Cells were collected at designated time points for further analysis. *4EBP1, Sufu, Rictor,* and *Sin1* KD cells were generated by transducing NIH 3T3 cells with lentiviruses expressing shRNAs. We produced replication-incompetent lentiviruses by transfecting 293T cells with helper vectors and shRNA vectors purchased from transOMIC technologies. The shRNA vectors used were TLHSU1400-1978 (*4EBP1*), TLMSU1400-24069 (*Sufu*), TLMSU1400-78757 (*Rictor*), and TLMSU1400-227743 (*Sin1*).

### Polysome purification

NIH 3T3 cells were incubated with CHX (10 μg/mL) for 15 min and lysed in hypotonic buffer (20 mM potassium acetate, 12 mM magnesium acetate, 20 mM Tris-HCl, pH 7.4) by Dounce homogenization (35 strokes). Cell debris and nuclei were removed by centrifugation at 14,000×g for 10 min. Supernatants were layered on 1 mL of a 17%–47% (wt/vol) sucrose gradient (10 mM sodium chloride, 12 mM magnesium chloride, 20 mM Tris-HCl, pH 7.4) and centrifuged for 1.5 h at 35,000 rpm in a TLS-55 Beckman rotor. Ten fractions were manually collected from each sample by pipetting, and the fractionated samples were concentrated with Vivaspin centrifugal concentrators (Sartorius, Elk Grove, IL) for further analysis.

### Purification and culture of GNP and medulloblastoma cells

To purify GNPs and medulloblastoma cells, cerebella of P7 mice were dissociated with Accutase (ThermoFisher, catalog no. 11110501), resuspended in Neurobasal medium (ThermoFisher, catalog no. 21103-049), and separated by Percoll density-gradient (60%–30%) centrifugation (3000 rpm for 15 min). Purified GNPs or medulloblastoma cells were grown in culture in Neurobasal medium containing 1× PSG. GNPs were treated with SAG (200nM; Cayman Chemical, catalog no. 11914), Torin1 (400 nM), or INK128 (400 nM) as indicated in the text or figures. Cells were collected at designated time points for further analysis.

### Mice

We used the following mouse strains: *SmoM2*^*loxP*^ (Jackson Laboratory, JAX stock # 005130), *GFAP::Cre* (JAX stock # 004600), *Raptor*^*loxP*^ (JAX stock # 013188), *Rictor*^*loxP*^ (JAX stock # 020649), and *Ptch1*^*loxP*^ (JAX stock # 012457). All mice were maintained on a mixed genetic background. We crossed females carrying the *loxP*-flanked alleles and males carrying the *Cre* alleles to induce Cre-mediated recombination in the brains of offspring. All mice were maintained on a 12 h dark/light cycle and housed with a maximum of five mice of the same sex per cage. We used animals of both sexes for the experiments. All animal procedures were approved by the Institutional Animal Care and Use Committee of St Jude Children’s Research Hospital.

### Oral gavage

INK128 (Selleck Chemicals, catalog no. S2811) was dissolved to 20 mg/mL in vehicle containing 5% n-methylpyrrolidone (Sigma-Aldrich), 15% polyvinylpyrrolidone K 30 (Sigma-Aldrich), and 80% water. INK128 (0.1, 0.15, or 0.2 mg/kg BW) or vehicle (2.5 μL/g BW) was administered daily by oral gavage from P10 until the designated time point. For survival analyses, we administered INK128 (0.2 mg/kg BW) or vehicle to mice daily from P7 and euthanized the animals when they became moribund.

### Immunoblotting

Cells were disrupted with RIPA lysis buffer (50 mM Tris-HCl pH 7.4, 100 mM NaCl, 5 mM EDTA, 1% Triton X-100) containing protease and phosphatase inhibitors (Roche), incubated on ice for 15 min, and centrifuged for 10 min at 14,000×g at 4°C. For brain samples, 100 μg of tissue was dissected and homogenized in RIPA buffer with a motor tissue grinder (Fisher Scientific, catalog no. 12-141-361). Tissue debris was pelleted and removed by centrifugation for 10 min at 14,000×g at 4°C. The protein concentrations in the supernatants were determined by DC Protein Assay (Bio-Rad, catalog no. 5000112). Supernatants containing 3 to 5 μg of protein were subject to SDS-PAGE and analyzed using antibodies to the following: p-4EBP1 T37/T46 (Cell Signaling Technology, catalog no. 2855), 4EBP1 (Cell Signaling Technology, catalog no. 9644), β-actin (Sigma-Aldrich, catalog no. A5441), p-Akt S473 (Cell Signaling Technology, catalog no. 4060), Akt (Cell Signaling Technology, catalog no. 2966), EIF4E (Cell Signaling Technology, catalog no. 2067), GLI1 (Cell Signaling Technology, catalog no. 2534), IFT88 (Proteintech, catalog no. 13967-1-AP), Ki67 (Abcam, catalog no. ab16667), MTOR (Cell Signaling Technology, catalog no. 2983), PCNA (Santa Cruz Biotechnology, catalog no. sc56), Raptor (Cell Signaling Technology, catalog no. 2280), RICTOR (Cell Signaling Technology, catalog no. 2114), RPL7A (Cell Signaling Technology, catalog no. 2415), p-RPS6 S235 (Cell Signaling Technology, catalog no. 2211), RPS6 (Cell Signaling Technology, catalog no. 3944), SIN1 (Millipore, catalog no. 05-1044), SMO (Santa Cruz Biotechnology, catalog no. sc166685), ubiquitin (Santa Cruz Biotechnology, catalog no. sc-8017), and GFP/YFP (Aves Labs, catalog no. GFP1020). Each experiment was repeated at least three times, and representative results are shown in the figures.

### qRT-PCR

Total RNA was extracted using RNeasy Mini Kits (Qiagen). An aliquot of 200 ng of total RNA from each sample was reverse transcribed with Superscript III reverse transcriptase (ThermoFisher). Quantitative PCR analysis using SYBR green (ThermoFisher) was performed on an Applied Biosystems 7900 Real-Time PCR System. Transcript levels were normalized to the expression levels of *Actb* or *Rn18s*. The primer sequences used are shown in Supplementary Table 1.

### Immunostaining

Mice were perfused with 4% paraformaldehyde (PFA) in PBS. Their brains were dissected out, fixed overnight in 4% PFA at 4°C, washed in PBS overnight, cryoprotected in 30% sucrose, embedded in OCT, and sectioned at a slice thickness of 12 μm. Cells grown in culture were fixed with 4% PFA in PBS for 15 min at 4°C then washed with PBS. Brain sections or fixed cells were incubated with primary antibodies overnight at 4°C then incubated with secondary antibodies at room temperature for 2 h and stained with DAPI (10 μg/mL, Sigma) for 10 min. Coverslips were mounted on slides with Aqua-Poly/Mount (Polysciences). Images were acquired with a Zeiss AxioImager upright microscope or a Zeiss780 microscope and processed using Adobe Photoshop. We used the following antibodies: anti-acetylated tubulin antibody (Sigma, catalog no. T6973, diluted 1:1000), anti-Smo (Santa Cruz Biotechnology, catalog no. sc166685, diluted 1:50), anti-p-4EBP1 (Cell Signaling Technology, catalog no. 2855, diluted 1:200), and Alexa Fluor®–conjugated secondary antibodies (Invitrogen).

### Human tissue microarrays and Immunohistochemistry

Molecular subgrouping of medulloblastoma tissue was performed using immunohistochemical staining essentially as previously described but with some slight modification (Ellison et al., 2011). Antibodies to YAP1 (Santa Cruz Biotechnology clone 63.7, diluted 1:50), β-catenin (Ventana Medical Systems clone 14, no dilution required), and GAB1 (Santa Cruz Biotechnology clone H-7, diluted 1:400) were used with appropriate secondary reagents. Detection of phosphorylated-4EBP1 was performed using a monoclonal antibody (Cell Signaling Technology clone 236B4, diluted 1:800). Scoring of phospho-4EBP1 immunolabeling was performed as follows: each tumor was evaluated in a tissue microarray for positive or negative staining, and assigned a qualitative score based on whether there was low, focal staining (staining in less than 50% of the tumor cells), intermediate staining (weak staining in more than 50% of the cells), or high staining (high-intensity staining in more than 50% of the cells).

### Statistics

For the results of Western blot and qPCR analysis, at least three sample sets for each treatment were collected for analysis by the two-tailed, unpaired Student’s *t*-test. We used chi-square and Kruskal-Wallis tests, respectively, to compare the proportion of tumors stained for p-4EBP1 and the intensity of p-4EBP1 staining in each medulloblastoma subgroup. *P*-values of less than 0.05 were considered significant. No data points or animals were excluded from the analysis.

## AUTHOR CONTRIBUTIONS

C.-C.W. performed most of the experiments. S.H. contributed the *in vivo* analyses. B.A.O. analyzed the p-4EBP1 levels in human medulloblastomas. Y.H.Y. quantified cilia containing SMO. F.R. performed the *Sin1* and *Rictor* KD experiments. C.G.E. provided human medulloblastoma samples. C.C.W. and Y.-G.H. analyzed the data and wrote the manuscript with input from their co-authors. Y.-G.H. supervised the project.

## ACKNOWLEDGMENTS

We thank S. Baker, X. Cao, H. Chi, M.-J. Han, H.J. Kim, K.A. Laycock, and S. Ogden for comments on the manuscript. Y.-G.H. is supported by NIH/NCI Cancer Center Core Support grant CA021765 (to St. Jude Children’s Research Hospital), the Sontag Foundation Distinguished Scientist Award, a Whitehall Foundation Research Grant, and ALSAC.

## FIGURE LEGENDS

**Supplementary Figure S1. mTORC1 is required for HH signaling.**

(A) Simplified schematic of MTOR pathway showing reagents used to block MTOR, PI3K, or AKT. (B) Knockdown of SUFU induces GLI1 levels in the absence of SHH. (C) Torin1 and SANT1 completely blocked the *Gli1* mRNA increase induced by SAG but not that induced by FGF2.

**Supplementary Figure S2. MTOR does not affect SMO accumulation in cilia.**

(A) Immunofluorescence labeling showing the accumulation of SMO (green) in cilia (red) 2 h after SAG treatment in the presence or absence of Torin1. Cilia were labeled with an antibody against acetylated tubulin (AcTub). The boxed areas are examples of cilia that are enlarged in the lower panels for separate and merged channels. Scale bars = 5 μm (upper panels) and 1.25 μm (lower panels). (B) Quantification of the proportion of cilia containing SMO. * *P<*0.05; *n.s*. *P>* 0.05.

**Supplementary Figure S3. SHH increases SMO protein levels independently of transcription or protein stability.**

(A) SHH, but not EGF or FGF2, increased SMO protein levels in NIH 3T3 cells 24 h after treatment. FGF2 also increased GLI1 levels. (B) Knockdown of cilia with shRNA against *Ift88* (*shIft88*) suppressed the SAG-induced increase in *Gli1* mRNA. (C) *Smo* mRNA levels were not affected by SAG or KD of cilia. The levels show fold changes relative to those in untreated control cells. The results are the average ± SD of four independent experiments, each performed in triplicate. ** *P<* 0.01. (D) Inhibitors of proteasomes (MG132, bortezomib) or lysosomes (chloroquine) did not increase the SMO protein levels. The accumulation of ubiquitinated proteins confirmed the disruption of the proteasomes. (E) Inhibition of the proteasomes by MG132 did not rescue the Torin1-mediated suppression of the SMO protein levels in NIH 3T3 cells co-treated with SAG and Torin1.

**Supplementary Figure S4. p-4EBP1 levels are high in SHH-subgroup medulloblastoma.** Representative images of p-4EBP1 staining in human medulloblastomas. For each human tumor in a tissue microarray, the staining intensity was scored from 1 to 4: no staining (A, scored as 0), low staining (B, scored as 1), intermediate staining (C, scored as 2), and high staining (D, scored as 3). Scale bars = 200 μm.

